# In situ polyadenylation enables spatial mapping of the total transcriptome

**DOI:** 10.1101/2022.04.20.488964

**Authors:** David W. McKellar, Madhav Mantri, Meleana Hinchman, John S.L. Parker, Praveen Sethupathy, Benjamin D. Cosgrove, Iwijn De Vlaminck

## Abstract

Spatial transcriptomics reveals the spatial context of gene expression, but current methods are limited to assaying polyadenylated (A-tailed) RNA transcripts. Here we demonstrate that enzymatic in situ polyadenylation of RNA enables detection of the full spectrum of RNAs, expanding the scope of sequencing-based spatial transcriptomics to the total transcriptome. We apply this Spatial Total RNA-Sequencing (STRS) approach to study skeletal muscle regeneration and viral-induced myocarditis. Our analyses reveal the spatial patterns of noncoding RNA expression with near-cellular resolution, identify spatially defined expression of noncoding transcripts in skeletal muscle regeneration, and highlight host transcriptional responses associated with local viral RNA abundance. In situ polyadenylation requires the addition of only a single step to a widely used protocol for spatial RNA-sequencing, and thus could be broadly and quickly adopted. Spatial RNA-sequencing of the total transcriptome will enable new insights into spatial gene regulation and biology.

## INTRODUCTION

Spatial transcriptomics reveals the spatial context of gene expression, but current methods are limited to assaying polyadenylated (A-tailed) RNA transcripts. Here we demonstrate that enzymatic *in situ* polyadenylation of RNA enables detection of the full spectrum of RNAs, expanding the scope of sequencing-based spatial transcriptomics to the total transcriptome. We apply this Spatial Total RNA-Sequencing (STRS) approach to study skeletal muscle regeneration and viral-induced myocarditis. Our analyses reveal the spatial patterns of noncoding RNA expression with near-cellular resolution, identify spatially defined expression of noncoding transcripts in skeletal muscle regeneration, and highlight host transcriptional responses associated with local viral RNA abundance. *In situ* polyadenylation requires the addition of only a single step to a widely used protocol for spatial RNA-sequencing, and thus could be broadly and quickly adopted. Spatial RNA-sequencing of the total transcriptome will enable new insights into spatial gene regulation and biology.

## MAIN TEXT

Spatial transcriptomics provides insight into the spatial context of gene expression^1–5^. Current methods are restricted to capturing polyadenylated transcripts and are not sensitive to many species of non-A-tailed RNAs, including microRNAs, newly transcribed RNAs, and many non-host RNAs. Extending the scope of spatial transcriptomics to the total transcriptome would enable observation of spatial distributions of regulatory RNAs and their targets, link non-host RNAs and host transcriptional responses, and deepen our understanding of spatial biology.

Here, we demonstrate Spatial Total RNA-Sequencing (STRS), a method that enables spatial profiling of both the A-tailed and non-A-tailed transcriptome. This is achieved with a simple modification of a commercially available protocol for spatial RNA-sequencing. STRS uses poly(A) polymerase to add poly(A) tails to RNAs *in situ*. STRS otherwise follows conventional protocols to capture, spatially barcode, and sequence RNAs. STRS is compatible with existing approaches for sequencing-based spatial transcriptomics, is straightforward to implement, and adds minimal cost and time to an already widely used commercially available workflow. STRS enables the capture of many RNAs that are missed by conventional workflows, including noncoding RNAs, newly transcribed RNAs, and viral RNAs. To demonstrate the versatility of the method, we applied STRS to study the regeneration of skeletal muscle after injury and the pathogenesis of viral-induced myocarditis.

## RESULTS

### In situ polyadenylation enables capture of coding and noncoding RNAs

STRS adds a single step to a commercially available method for spatial RNA-sequencing (Visium Spatial Gene Expression, 10x Genomics) to capture the total transcriptome^6^. As in the Visium method, the sample is first sectioned, fixed with methanol, and stained for histology. After imaging, the sample is rehydrated and then incubated with yeast poly(A) polymerase for 25 minutes at 37°C. This enzyme adds poly(A) tails to the 3’ end of all RNAs so that endogenous poly(A) tails are extended and non-A-tailed transcripts are polyadenylated. After *in situ* polyadenylation, STRS again follows the Visium protocol without modification (**Fig 1a**). One important feature of the Visium method that we leverage in STRS, is its use of a strand-aware library preparation. We found that strandedness is critical for the study of noncoding and antisense RNAs (see below) and must be considered in bioinformatic analyses (**Fig S1**).

**Figure 1.**
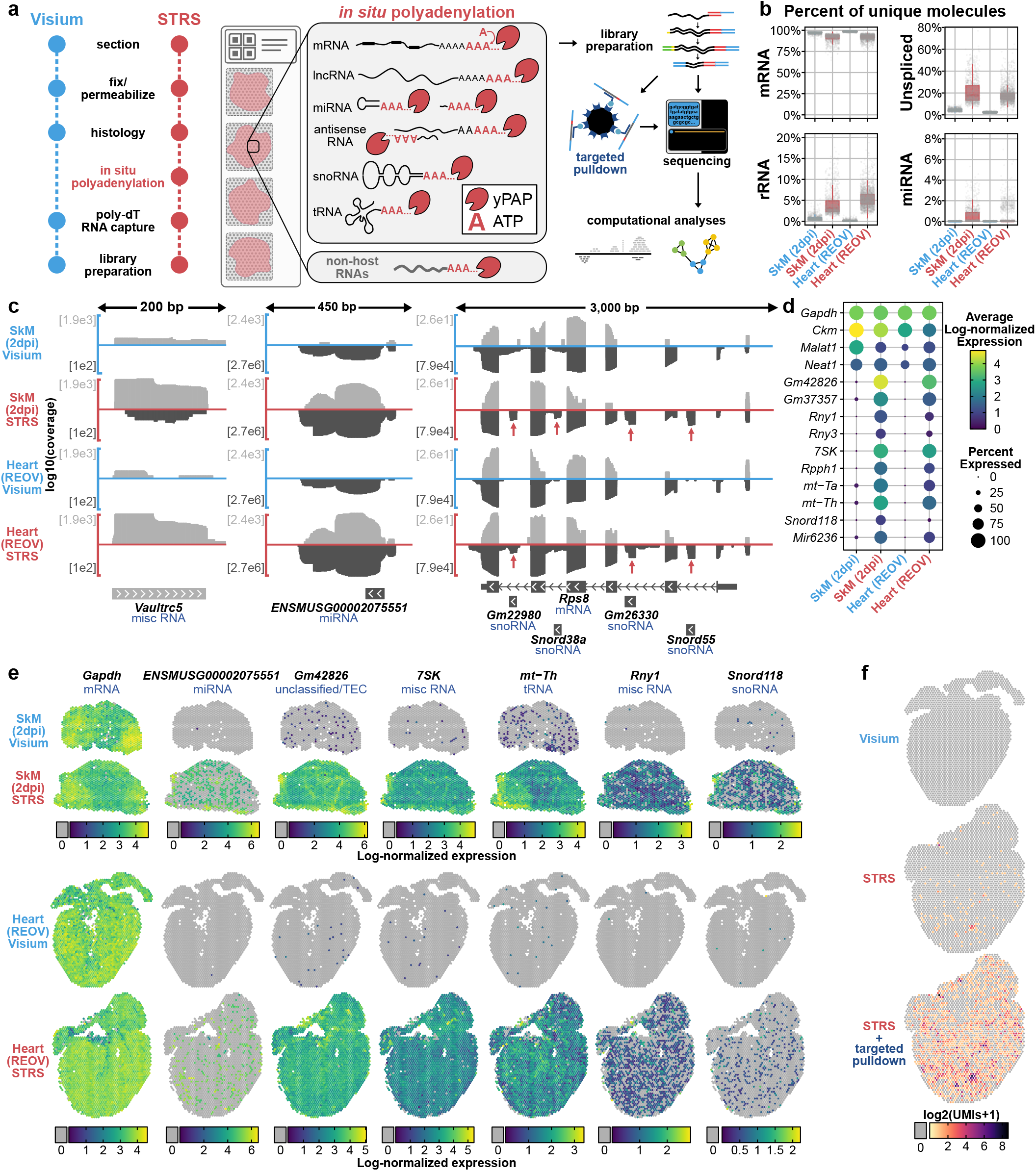
*In situ* polyadenylation enables spatial profiling of noncoding and non-host RNAs. **(a)** Workflow for Spatial Total RNA-Sequencing (STRS). **(b)** Comparison of select RNA biotypes between Visium and STRS datasets. Y-axis shows the percent of unique molecules (UMIs) for each spot. **(c)** Detection of coding and noncoding RNAs between Visium and STRS workflows. Color scale shows average log-normalized UMI counts. Dot size shows the percent of spots in which each RNA was detected. **(d)** Log10-transformed coverage of deduplicated reads mapping to sense (light gray) and antisense (dark gray) strands at the *Vaultrc5, ENSMUSG00002075551*, and *Rps8* loci. Annotations shown are from GENCODE M28 and include one of the five isoforms for *Rps8* as well as the four intragenic features within introns of *Rps8*. **(e)** Spatial maps of coding and noncoding transcripts for Visium and STRS workflows. Spots in which the transcript was not detected are shown as gray. **(f)** Detection of reovirus transcripts using the standard workflow, STRS, and STRS with targeted pulldown enrichment. Spots in which the virus was not detected are shown as gray.

To test the performance and versatility of STRS, we applied it to two distinct mouse tissue types: injured hindlimb muscle^5^ and virally infected heart tissue^4^. We quantified the percentage of unique molecules (UMIs) as a function of RNA biotype (GENCODE M28 annotations; **Fig 1b**). Compared to the Visium method, we found similar counts for protein coding and other endogenously polyadenylated transcripts (**Fig S1-2**). STRS enabled robust detection of several types of noncoding RNAs which are poorly recovered or not detected at all by the Visium method, including ribosomal RNAs (rRNAs; mean of 5.4% and 2.6% of UMIs for STRS and Visium respectively; computed across all Visium and STRS samples included in this study), microRNAs (miRNAs; 0.4% in STRS versus 0.004% in Visium), transfer RNAs (tRNAs; 0.4% in STRS versus 0.02% in Visium), small nucleolar RNAs (snoRNAs; 0.2% in STRS versus 0.002% in Visium), and several other biotypes (**Fig 1b, Fig S3-4**). STRS libraries also had an increased fraction of unspliced transcripts (2.7% in Visium versus 18.3% in STRS). Unspliced or nascent RNA counts have been used to predict transcriptional trajectories for single cells. Improved detection of nascent RNAs may enable more accurate trajectory imputation and reveal the dynamics of spatial gene expression. Finally, STRS libraries had an increased fraction of reads which map to intergenic regions, reflecting increased capture of unannotated transcriptional products (22.2% in STRS versus 9.5% in Visium; **Fig S1b-c**). We found that STRS captured many RNAs which were not present in Visium libraries. Many of these features map outside of or antisense to known annotations (**Fig 1c**). We also found that STRS detected many noncoding transcripts which are intragenic to other genes (**Fig 1c**). Standard short-read sequencing was sufficient to delineate these features from the surrounding host genes, as reflected by the expression count matrices for STRS versus Visium data (**Fig 1d**). Most importantly, we were able to spatially map each of these features and visualize spatial patterns of gene expression (**Fig 1e**). We found that features which were incompletely annotated (*ENSMUSG00002075551*) showed sparse spatial expression. Several highly abundant genes showed homogenous patterns of expression, reflecting putative (*Gm42826*) or known (*7SK*) housekeeping roles^7^.

We also asked whether *in situ* polyadenylation enables capture of non-A-tailed viral RNA. To this end, we assayed murine heart tissues infected with Type 1-Lang reovirus (REOV), a segmented double-stranded RNA virus that expresses ten transcripts which are not polyadenylated. No reovirus transcripts were detected with the Visium workflow, whereas STRS enabled detection of more than 200 UMIs representing all ten reovirus gene segments (**Fig 1f**). To deeply profile viral RNAs, we performed targeted enrichment of viral-derived cDNA from the final sequencing libraries and re-sequenced the products. This enrichment led to a further ∼26-fold increase of the mean viral UMIs captured per spot (minimum L1 segment with 262 UMIs, maximum S4 segment with 1095 UMIs). Taken together, these findings demonstrate that STRS enables the study of many types of RNAs that are not detectable with existing technologies.

### Spatial total RNA-sequencing reveals spatial patterns of gene regulation in skeletal muscle regeneration

Skeletal muscle regeneration is a coordinated system guided by complex gene regulatory networks^5,8–12^. We applied STRS to spatially map the coding and noncoding transcriptome in a mouse model of skeletal muscle regeneration. We injured tibialis anterior muscles and then collected tissues at 2-, 5-, and 7-days post-injury (dpi) in addition to an uninjured control (**Methods**). H&E imaging showed immune infiltration centrally within tissue sections at 2 and 5dpi, which was mostly resolved by 7dpi (**Fig 2a**). Unsupervised clustering identified spots in the injury loci, spots around the border of the injury loci, and spots under intact myofibers (**Fig 2b, Methods**).

**Figure 2.**
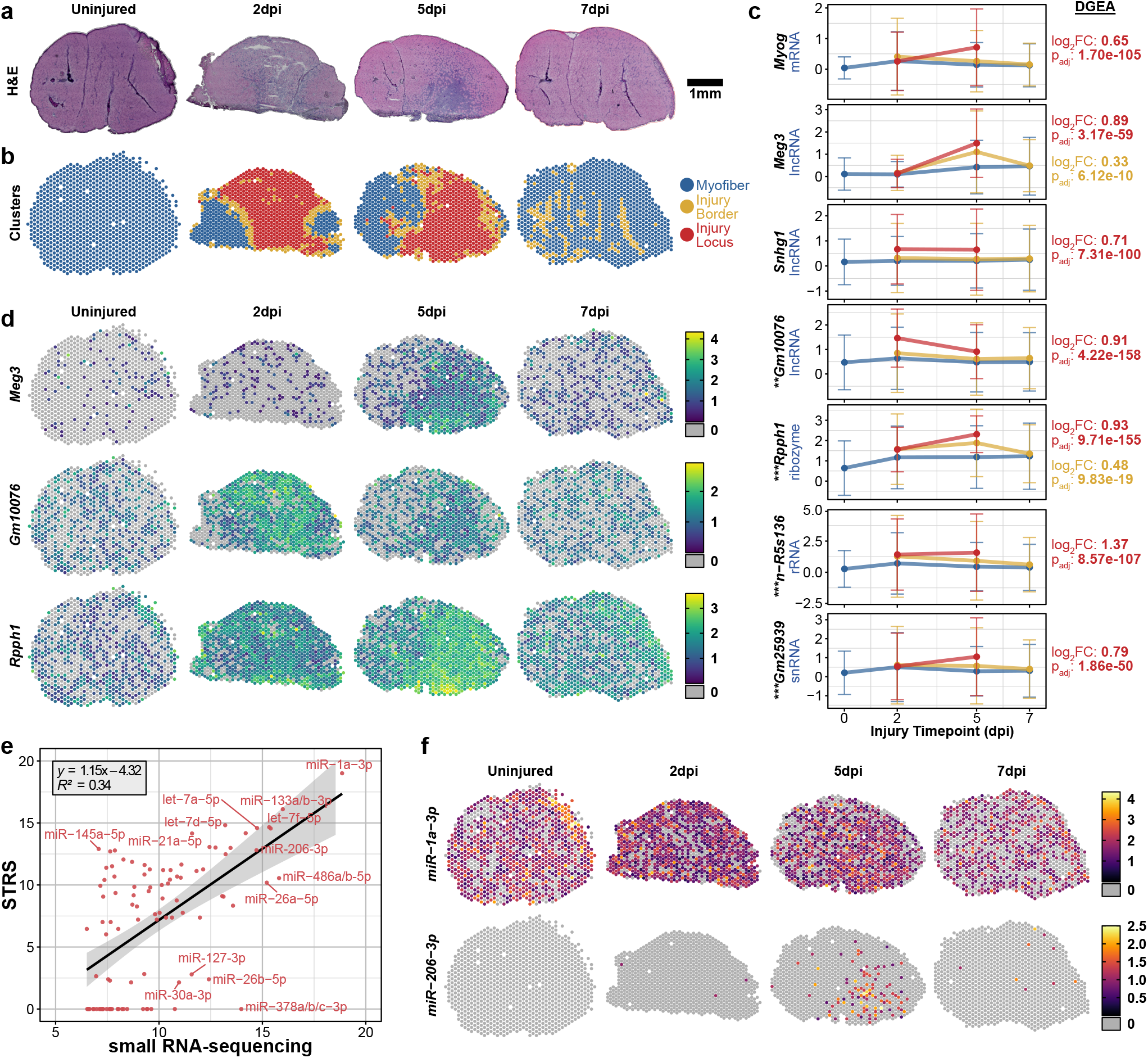
Spatial total RNA-sequencing of regenerating skeletal muscle **(a)** H&E histology of mouse tibialis anterior muscles collected 2-, 5-, and 7-days post-injury (dpi). **(b)** Clustering of spot transcriptomes based on total transcriptome repertoires (see **Methods**). **(c)** Differentially expressed RNAs across regional clusters. Y-axis shows log-normalized expression of each feature. Mean expression across each cluster is reported, colored according to the legend in (b). Error bars show standard deviation. Reported statistics to the right of plots reflect differential gene expression analysis performed across clusters on merged STRS samples (Wilcoxon, see Methods). Asterisks next to transcript names reflect differential expression analysis performed across skeletal muscle Visium and STRS samples (**p_val_adj<10^−50^, ***p_val_adj<10^−150^; Wilcoxon, see Methods) **(d)** Spatial maps for select features from (c). **(e)** Mature miRNA expression detected by STRS. Color scale shows log-normalized miRNA counts, quantified by miRge3.0 (**Methods**). (**f**) Average detection of miRNAs compared between small RNA-sequencing (n=8) and STRS (n=4). Axes show log2 counts per million transcripts, normalized to the total number of transcripts which map to small RNA loci with miRge3.0. The top 100 most abundant miRNAs detected by small RNA-sequencing are shown.

We performed differential gene expression analysis across the regional clusters to identify noncoding RNAs specific to the injury locus (**Fig 2c, Methods**). We found several RNAs which were spatiotemporally associated with injury locus, many of which are undetected or poorly detected by Visium (**Fig 2c-d**). *Meg3* is an endogenously polyadenylated lncRNA which has been shown to regulate myoblast differentiation in vitro. We found *Meg3* expression to be confined to the injury locus at 5dpi, when myoblast differentiation and myocyte fusion occurs^5,13^. *Gm10076*, a transcript with a biotype annotation conflict (Ensembl: lncRNA; NCBI: pseudogene) and no known function, was highly and specifically expressed within the injury locus 2dpi. *Gm10076* expression was reduced but still localized to the injury site by 5dpi and returned to baseline levels by 7dpi. *Rpph1*, a ribozyme and component of the RNase P ribonucleoprotein which has also been shown to play roles in tRNA and lncRNA biogenesis^14,15^, showed broad expression by 2dpi which peaked and localized to the injury site at 5 dpi. We also found that STRS captured high levels of antisense transcripts for *Rpph1* which were not detected by the Visium chemistry. This demonstrates that STRS can robustly profile both polyadenylated and non-polyadenylated RNAs across heterogeneous tissues.

The role of miRNAs in skeletal muscle regeneration has been well-established^11,16–18^. Mature miRNAs^19^ are ∼22 nucleotides long, are not polyadenylated, and are not captured by the standard Visium workflow (**Fig S5**). We asked if STRS was able to detect mature miRNAs. We first generated matched bulk small RNAseq libraries from entire tibialis anterior muscles as a gold standard reference (n=2 per timepoint). We used miRge3.0^20^ to quantify mature miRNA abundance in the STRS and matched small RNA-sequencing libraries (**Methods**). We found strong correlation in the abundance of the most highly expressed miRNAs between STRS and small RNA-sequencing, with only minor drop-out of lowly expressed miRNAs (**Fig 2e, Fig S5**). We identified many examples of mature miRNA expression in STRS data, including expression of classic “myomiRs”, *miR-1a-3p, miR-133a/b-3p*, and *miR-206-3p* (**Fig 2f**)^21^. Consistent with previous studies^22^, we detected static expression of *miR-1a-3p* across all four timepoints (**Fig 2d**), whereas *miR-206-3p* was highly expressed within the injury locus five days post-injury, with very low levels of expression detected at other timepoints.

### Spatial total transcriptomics spatially resolves viral infection of the murine heart

We next explored the potential for STRS to profile host-virus interactions in a mouse model of viral-induced myocarditis. We orally infected neonatal mice with type 1-Lang reovirus (REOV), a double-stranded RNA virus with gene transcripts that are not polyadenylated. Within seven days of oral infection, REOV spreads to the heart and causes myocarditis^23–25^. We performed Visium and STRS on hearts collected from REOV-infected and saline-injected control mice (**Fig 3a**). We found that reovirus transcripts were only detected in the infected heart via STRS and that targeted enrichment of reovirus transcripts enabled deeper profiling of viral infection (**Fig 1d, 3a; Methods**). Mapping these reads across the tissue revealed pervasive infection across the heart (1,329/2,501 or 53% spots under the tissue; **Fig 1d**). Foci containing high viral UMI counts overlapped with myocarditic regions as identified by histology.

**Figure 3.**
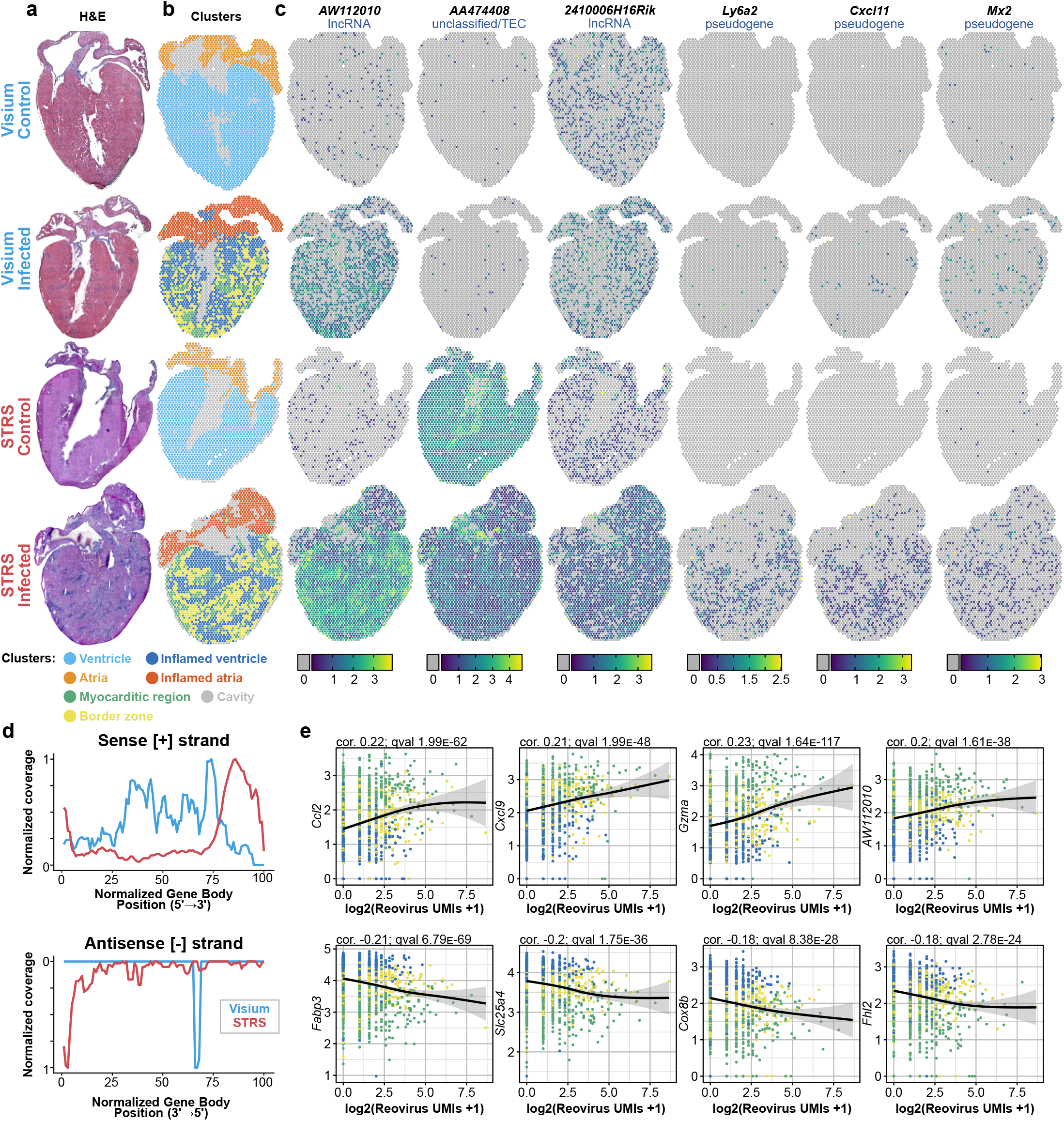
STRS enables simultaneous analysis of viral infection and host response. **(a)** H&E staining of control and reovirus-infected hearts, collected using the standard Visium workflow and STRS. **(b)** Tissue regions identified through unsupervised clustering of spot transcriptomes. **(c)** Log-normalized expression of noncoding and coding RNAs which are highly expressed in myocarditic regions. Spots in which transcripts were not detected are shown in gray. **(d)** Normalized coverage of deduplicated reads for the sense [+] and antisense [-] strands of all ten reovirus gene segments. X-axis shows the length-normalized position across the gene bodies of all ten reovirus segments. Note that the peak in antisense [-] coverage for the Visium sample (blue) corresponds to only 11 total reads. **(e)** Co-expression of pulldown-enriched reovirus UMIs versus infection-associated genes in spots underneath inflamed and myocarditic tissue. Spots are colored according to legend in (b). Correlation and q-value reported are from general additive model analysis (**Methods**).

We next compared the read coverage profiles across the ten REOV gene segments for REOV-enriched libraries from Visium and STRS samples (**Fig 3d**). As expected, STRS libraries had a peak in coverage at the 3’ end of viral gene segments. In contrast, the REOV-enriched Visium reads contained peaks in the middle of viral gene segments as expected for a chemistry that relies on the spurious capture of viral RNA at poly(A) repeats within the transcripts^26^. Interestingly, we found that STRS led to an overrepresentation of reads from the 5’ end of the sense [+] strand of all ten REOV segments. These reads may represent incomplete transcripts generated by transcriptional pausing of the REOV RNA polymerase or transcripts undergoing 3’ exonucleolytic degradation. Finally, we detected the 3’ end of the antisense [-] strand for nine of the ten segments of the reovirus genome, suggesting that STRS captures both strands of the dsRNA reovirus genome (**Fig 3d**). These antisense reads were present at an average ratio of ∼1:40 compared to the sense reads. The current model for synthesis of reovirus dsRNA posits that dsRNA synthesis only occurs within a viral core particle after packaging of the ten viral positive-sense RNAs. There are several possible explanations for our detection of the antisense strands. One is that we are detecting negative-strand viral RNA that is part of dsRNA that has been released from damaged viral particles either within the cytoplasm or within lysosomes. dsRNA released within endolysosomes can be transported into the cytoplasm by RNA transmembrane receptors SIDT1 and SIDT2^27,28^. A second possibility is that antisense [-] viral RNA is synthesized prior to packaging of dsRNA into viral particles.

Because STRS efficiently recovers viral RNA, we were able to directly correlate host transcriptomic responses with viral transcript counts for spots in inflamed regions (generalized additive model, **Methods**). We found inflammation-associated cytokine transcripts such as *Ccl2* and *Cxcl9*, and immune cell markers such as *Gzma* and *Trbc2* to be upregulated in spots with high viral counts (**Fig 3e**). We continued this analysis by performing unsupervised clustering (**Fig 3b**) and differential gene expression analysis to identify transcripts associated with infection which are more readily detected by STRS (**Fig 3c**). *AW112010*, which has recently been shown to regulate inflammatory T cell states, was only found in infected samples and was more abundant in the STRS data compared to Visium. We also found that STRS led to increased detection of putative protein-coding genes, including *Ly6a2, Cxcl11*, and *Mx2*, which were associated with infection. Interestingly, all three genes are annotated as pseudogenes in GENCODE annotations but have biotype conflicts with other databases. The increased abundance as measured by STRS could reflect differential mRNA polyadenylation for these transcripts. Overall, STRS enabled more robust, spatially mapped analysis of the host response to infection by increasing the breadth of captured transcript types and by providing direct comparison with viral transcript abundance.

## DISCUSSION

Here, we demonstrate *in situ* polyadenylation of RNA in sectioned tissue to enable Spatial Total RNA-Sequencing. Enzymatic polyadenylation is frequently implemented for bulk sequencing of total RNA and was recently adopted for single-cell RNA-sequencing^29,30^. STRS is the first implementation of *in situ* RNA-labeling for spatial RNA-sequencing. STRS has several notable features. First, STRS is compatible with a commercial workflow and requires the use of only one additional reagent. STRS can easily be adopted by others as it requires minimal additional experimental time (∼30 minutes) and cost (<$90 per sample) and does not require any specialized equipment. Second, because our RNA-labeling strategy is designed to work with poly(dT) reverse transcription, STRS is compatible with other sequencing-based spatial transcriptomics platforms. The resolution of our analyses is limited by the size and distribution of the barcoded spots on the Visium slides. Future iterations of STRS which use higher resolution RNA-capture platforms, including Slide-Seqv2^31^, BGI Stereo-seq^32^, or new versions of Visium, promise substantial improvements in spatial resolution. Because STRS is not targeted and does not require prior sequence information, it is easily adapted to new biological systems and is well-suited for assaying unknown RNAs, including novel RNAs or non-host transcripts.

We investigated the utility and versatility of STRS by applying it to two distinct models. First, we profiled the noncoding RNA repertoires of infiltrating immune cells and regenerating myogenic cells at injury loci in mouse muscle. Second, we analyzed the host transcriptome in response to mammalian orthoreovirus infection. Members of the Reoviridae family of viruses synthesize non-polyadenylated viral mRNAs as do Arenaviruses, and flaviviruses^33–35^. Because STRS can directly capture viral RNAs, we could directly compare viral RNA abundance to gene expression changes in heart tissue. This enabled identification of infection-related lncRNAs which were not detectable using standard techniques. Adding spatial context has clarified the underlying biology of gene expression measurements. STRS improves on these facets by extending the assayable transcriptome and enabling direct measurements of viral-derived RNA transcripts.

With STRS, we demonstrated the first method to simultaneously map miRNAs and the mRNAs on which they act. Because of their short length and known biases in adapter ligation, miRNAs are notoriously difficult to assay^36,37^. Furthermore, the Visium Gene Expression protocol uses a tagmentation-based library preparation which depletes short molecules by either cutting the UMI/spot barcode or by producing a read which is too short to confidently align to the genome. Despite these issues, we showed robust detection for several known myomiRs and strong correlation with a gold-standard bulk method that does not suffer from ligation or length biases. With future improvements to the library preparation strategy, many of these hurdles can be further reduced.

This work highlights opportunities for improvements in current bioinformatic tools and resources for single-cell and spatial transcriptomics. Current alignment and transcript counting tools are not optimized for total RNA data and genome annotations are incomplete outside of protein coding genes. Furthermore, new tools that go beyond UMI counts and better leverage the wealth of information in sequence read alignment patterns are likely to be highly impactful.

In conclusion, we have demonstrated a versatile strategy for spatial mapping of the total transcriptome. We think STRS will expand the scope of spatial transcriptomics and enable new types of analyses on spatial gene regulation at tissue scale.

## METHODS

### Mice

The Cornell University Institutional Animal Care and Use Committee (IACUC) approved all animal protocols, and experiments were performed incompliance with its institutional guidelines. For skeletal muscle samples, adult female C57BL/6J mice were obtained from Jackson Laboratories (#000664; Bar Harbor, ME) and were used at 6 months of age. For heart samples, confirmed pregnant female C57BL/6J mice were ordered from Jackson Laboratories to be delivered at embryonic stage E14.5.

### Viral infection

Litters weighing 3 gram/ pup were orally gavaged using intramedic tubing (Becton Dickinson Cat #427401 with 50 μl with 10^7^ PFU reovirus type 1-lang (T1L) strain in 1X phosphate buffered saline (PBS) containing green food color (McCormick) via a 1ml tuberculin slip tip syringe (BD 309659) and 30G x 1/2 needle (BD Cat #305106). Litters treated with 1X PBS containing green food color alone on the same day were used as mock controls for the respective infection groups. The mock-infected and reovirus-infected pups were monitored and weighed daily until the time points used in the study (7 days post infection). After dissection, samples were embedded in O.C.T. Compound (Tissue-Tek) and frozen fresh in liquid nitrogen.

### Muscle injury

To induce muscle injury, both tibialis anterior muscles of old (20 months) C57BL/6J mice were injected with 10µl of notexin (10 µg/ml; Latoxan; France). Either before injury or 2-, 5-, or 7-days post-injury (dpi), mice were sacrificed and tibialis anterior muscles were collected. After dissection, samples were embedded in O.C.T. Compound (Tissue-Tek) and frozen fresh in liquid nitrogen.

### In situ polyadenylation and spatial total RNA-sequencing (STRS)

Spatial total RNA-sequencing was performed using a modified version of the Visium protocol. 10um thick tissue sections were mounted onto the Visium Spatial Gene Expression v1 slides. For heart samples, one tissue section was placed into each 6×6mm capture area. For skeletal muscle samples, two tibialis anterior sections were placed into each capture area. After sectioning, tissue sections were fixed in methanol for 20 minutes at -20°C. Next, H&E staining was performed according to the Visium protocol, and tissue sections were imaged on a Zeiss Axio Observer Z1 Microscope using a Zeiss Axiocam 305 color camera. H&E images were shading corrected, stitched, rotated, thresholded, and exported as TIFF files using Zen 3.1 software (Blue edition). After imaging, the slide was placed into the Visium Slide Cassette. *In situ* polyadenylation was then performed using yeast Poly(A) Polymerase (yPAP; Thermo Scientific, Cat #74225Z25KU). First, samples were equilibrated by adding 100µl of 1X wash buffer (20µl 5X yPAP Reaction Buffer, 2µl 40U/µl Protector RNase Inhibitor, 78µl nuclease-free H2O) to each capture area and incubating at room temperature for 30 seconds. The buffer was then removed. Next, 75µl of yPAP enzyme mix (15µl 5X yPAP Reaction Buffer, 3µl of 600U/µl yPAP enzyme, 1.5µl 25mM ATP, 3µl 40U/µl Protector RNase Inhibitor, 52.5µl nuclease-free H2O) was added to each reaction chamber. STRS was also tested with 20U/µl of SUPERase-In RNase-Inhibitor, but we found that SUPERase was not able to prevent degradation of longer transcripts during *in situ* polyadenylation (**Fig S6c-d**). The reaction chambers were then sealed, and the slide cassette was incubated at 37°C for 25 minutes. The enzyme mix was then removed. Prior to running STRS, optimal tissue permeabilization time for both heart and skeletal muscle samples was determined to be 15 minutes using the Visium Tissue Optimization Kit from 10x Genomics. Following *in situ* polyadenylation, the standard Visium library preparation was followed to generate cDNA and final sequencing libraries. The libraries were then pooled and sequenced according to guidelines in the Visium Spatial Gene Expression protocol using either a NextSeq 500 or NextSeq 2000 (Illumina, San Diego, CA).

### Small RNA-sequencing

For skeletal muscle samples, following the injury time course, tibialis anterior muscles were dissected and snap frozen with liquid nitrogen. The Norgen Total RNA Purification Kit (Cat. 17200) was used to extract RNA from 10 mg of tissue for each sample. For heart samples, following the infection time course, hearts were dissected, embedded in OCT, and frozen in liquid nitrogen. RNA was extracted with Trizol (Invitrogen, Cat. 15596026) and glycogen precipitation for a small fraction of each of the heart samples. RNA quality was assessed via High Sensitivity RNA ScreenTape Analysis (Agilent, Cat. 5067-5579) and all samples had RNA integrity numbers greater than or equal to 7.

Small RNA sequencing was performed at the Genome Sequencing Facility of Greehey Children’s Cancer Research Institute at the University of Texas Health Science Center at San Antonio. Libraries were prepared using the TriLink CleanTag Small RNA Ligation kit (TriLink Biotechnologies, San Diego, CA). Libraries were sequenced with single-end 50× using a HiSeq2500 (Illumina, San Diego, CA).

### Preprocessing and alignment of Spatial Total RNA-Sequencing data

All code used to process and analyze these data can be found at https://github.com/mckellardw/STRS. An outline of the pipelines used for preprocessing and alignment is shown in **Fig S1**.

Reads were first trimmed using cutadapt v3.4^38^ to remove the following sequences: 1) poly(A) sequences from the three prime ends of reads, 2) the template switch oligonucleotide sequence from the five prime end of reads which are derived from the Visium Gene Expression kit (sequence: CCCATGTACTCTGCGTTGATACCACTGCTT), 3) poly(G) artifacts from the three prime ends of reads, which are produced by the Illumina two-color sequencing chemistry when cDNA molecules are shorter than the final read length, and 4) the reverse complement of the template switching oligonucleotide sequence from the five prime ends of reads (sequence: AAGCAGTGGTATCAACGCAGAGTACATGGG). Next, reads were aligned using either STAR v2.7.10a^39^ or kallisto v0.48.0^40^. Workflows were written using Snakemake v6.1.0^41^.

For STAR, the genomic reference was generated from the GRCm39 reference sequence using GENCODE M28 annotations. For STAR alignment, the following parameters, based on work by Isakova et al, were used: outFilterMismatchNoverLmax=0.05, outFilterMatchNmin=16, outFilterScoreMinOverLread=0, outFilterMatchNminOverLread=0, outFilterMultimapNmax=50. Aligned reads were deduplicated for visualization using umi-tools v1.1.2^42^.

For kallisto, a transcriptomic reference was also generated using the GRCm39 reference sequence and GENCODE M28 annotations. The default k-mer length of 31 was used to generate the kallisto reference. Reads were pseudoaligned using the ‘kallisto bus’ command with the chemistry set to “VISIUM” and the ‘fr-stranded’ flag activated to enable strand-aware quantification. Pseudoaligned reads were then quantified using bustools v0.41.0. First, spot barcodes were corrected with ‘bustools correct’ using the “Visium-v1” whitelist provided in the Space Ranger software from 10x Genomics. Next, the output bus file was sorted and counted using ‘bustools sort’ and ‘bustools count’, respectively. To estimate the number of spliced and unspliced transcripts, reads pseudoaligned using kb-python v0.26.0, using the “lemanno” workflow.

Spots were manually selected based on the H&E images using Loupe Browser from 10x Genomics. Spatial locations for each spot were assigned using the Visium coordinates provided for each spot barcode by 10x Genomics in the Space Ranger software (“Visium-v1_coordinates.txt”). Downstream analyses with the output count matrices were then performed using Seurat v4.0.4^43,44^. In addition to manual selection, spots containing fewer than 500 detected features or fewer than 1000 unique molecules were removed from the analysis. Counts from multimapping features were collapsed into a single feature to simplify quantification.

### Mature microRNA quantification

For STRS data: after trimming (see above), barcode correction with STAR v2.7.10a, and UMI-aware deduplication with umi-tools v1.1.2, reads were split across all 4992 spot barcodes and analyzed using miRge3.0 v0.0.9^20^. Reads were aligned to the miRbase reference provided by the miRge3.0 authors. MiRNA counts were log-normalized according to the total number of counts detected by kallisto and scaled using a scaling factor of 1000. For small RNAseq data: Reads were first trimmed using trim_galore v0.6.5. Reads were then aligned and counted using miRge3.0 v0.0.9.

### Unsupervised clustering and differential gene expression analysis of spot transcriptomes

Spot UMI counts as generated by kallisto were used. First, counts were log-normalized and scaled using default parameters with Seurat. Principal component analysis was then performed on the top 2000 most variable features for each tissue slice individually. Finally, unsupervised clustering was performed using the ‘FindClusters()’ function from Seurat. The top principal components which accounted for 95% of variance within the data were used for clustering. For skeletal muscle samples, a clustering resolution was set to 0.8. For heart samples, clustering resolution was set to 1.0. Default options were used for all other parameters. Finally, clusters were merged according to similar gene expression patterns and based on histology of the tissue under each subcluster.

Differential gene expression analysis was performed using the ‘FindAllMarkers()’ function from Seurat. Default parameters were used, including the use of the Wilcoxon ranked sum test to identify differentially expressed genes. To identify features enriched in the skeletal muscle STRS datasets, all Visium and STRS were first merged and compared according to the method used (Visium vs. STRS). To identify cluster-specific gene expression patterns, skeletal muscle samples were first clustered as described above individually. STRS samples were then merged, and differential gene expression analysis was performed across the three injury region groups.

### Targeted pulldown enrichment of viral fragments

We performed hybridization-based enrichment of viral fragments on the Visium and STRS libraries for reovirus-infected hearts using the xGen Hybridization and Wash Kit (IDT; 1080577)^4^. In this approach, a panel of 5’-biotinylated oligonucleotides was used for capture and pulldown of target molecules of interest, which were then PCR amplified and sequenced. We designed a panel of 202 biotinylated probes tiled across the entire reovirus T1L genome to selectively sequence viral molecules from the sequencing libraries (**Table S1**). After fragmentation and indexing of cDNA, 300ng of the final Visium or STRS sequencing libraries from reovirus-infected hearts were used for xGen hybridization capture using the xGen NGS Target Enrichment Kit protocol provided by the manufacturer. One round of hybridization capture was performed for the STRS library followed by 14 cycles of PCR amplification. Because of the reduced number of captured molecules, two rounds of hybridization were performed on the Visium libraries. Enriched VIsium libraries were PCR-amplified for 18 cycles after the first round of hybridization and by 5 cycles after the second round of hybridization. Post-enrichment products were pooled and sequenced on the Illumina NextSeq 500.

### Correlation analysis between reovirus counts and host gene expression

We used a generative additive model (GAM) implemented in Monocle v2.18.0^45^ to find genes that vary with viral UMI count. A Seurat object for STRS data and viral UMI counts from the reovirus-infected heart was converted to a CellDataSet object using the ‘as.CellDataSet()’ command implemented in Seurat. The expression family was set to “negative binomial” as suggested for UMI count data in the Monocle documentation. The CellDataSet object was then preprocessed to estimate size factors and dispersion for all genes. Genes expressed in fewer than 10 spots were removed. Within the remaining genes, we then used the GAM implemented in the ‘differentialGeneTest()’ command in Monocle to identify genes that vary with log-transformed viral UMI counts. To find the direction in which these genes varied with viral UMI counts, we calculated the Pearson correlation for all genes with log2-transformed viral UMI counts.

## DATA AND CODE AVAILABILITY

Previously published spatial RNA-sequencing data were downloaded from Gene Expression Omnibus (GEO) and are available under the following accession numbers; regenerating skeletal muscle^5^ GSE161318, infected heart tissue^4^ GSE189636. Spatial Total RNA-Sequencing data generated in this study can be found on GEO under the accession number GSE200481. Small RNA-sequencing data are available on GEO under the accession number GSE200480 A detailed protocol for performing STRS as well as custom analysis scripts for aligning and processing STRS data can be found at https://github.com/mckellardw/STRS.

## ACKNOWLEDGMENTS

We thank Peter Schweitzer and colleagues in the Cornell Biotechnology Resource Center for their help with sequencing the libraries. We thank the Cornell Center for Animal Resources and Education for animal housing and care. We thank Ern Hwei Hannah Fong for helping with mouse procedures. We thank Michael Shanahan and Zhao Lai for their help in generating the small RNA-sequencing data. We thank Benjamin Grodner, Hao Shi, and other members of the Cosgrove and De Vlaminck labs for helpful discussions and feedback. This work was supported by the US National Institutes of Health (NIH) grants 1DP2AI138242 to IDV, R21AI144557 to IDV and JSP, NIH grant R01AG058630 to BDC and IDV, American Diabetes Association Pathway to Stop Diabetes Award 1-16-ACE-47 to PS, and T32EB023860 to DWM. The content is solely the responsibility of the authors and does not necessarily represent the official views of the NIH.

## AUTHOR CONTRIBUTIONS

DWM, IDV, and BDC designed the study. DWM, MM, and MH carried out the experiments. DWM and MM analyzed the data. DWM, MM, IDV, and BDC wrote the manuscript. All authors provided feedback and comments.

