## Supplemental Figures for "In situ polyadenylation enables spatial mapping of the total transcriptome"

Code repository: https://github.com/mckellardw/STRS

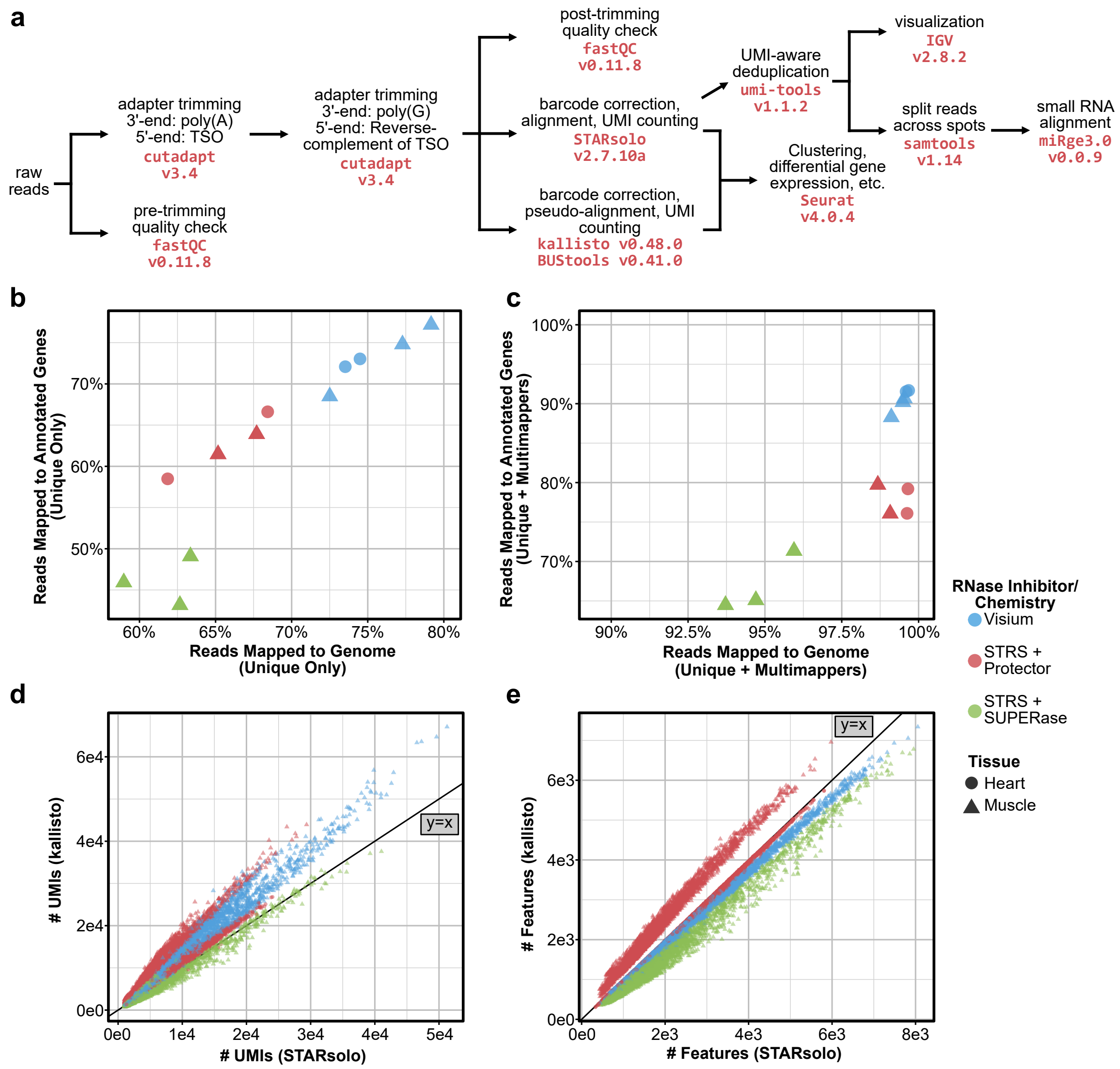


**Figure S1.** Comparison of bioinformatic analyses for Visium and Spatial Total RNA-Sequencing (STRS). (a) Bioinformatic tools and workflows used to preprocess, align, and quantitate transcripts. (b) STAR alignment rate for reads mapping to unique genomic position (x-axis) versus reads uniquely mapping to annotated regions (GENCODE M28 annotations) of the genome (y-axis). Each point represents an entire Visium capture area. Points are colored by sample preparation method (see Methods) and are shaped according to tissue type. (c) STAR alignment rate for reads mapping to unique or multiple positions along the genome (x-axis) versus reads uniquely mapping or multimapping to annotated regions (GENCODE M28 annotations) of the genome (y-axis). Spots are colored as in (b). (d) Number of unique molecules (UMIs) detected by STARsolo (x-axis) versus kallisto (y-axis). Each point represents a barcoded spot. Points are colored by sample preparation method (see Methods) and are shaped according to tissue type. (e) Number of features detected by STARsolo (x-axis) versus kallisto (y-axis). Spots are colored as in (d).

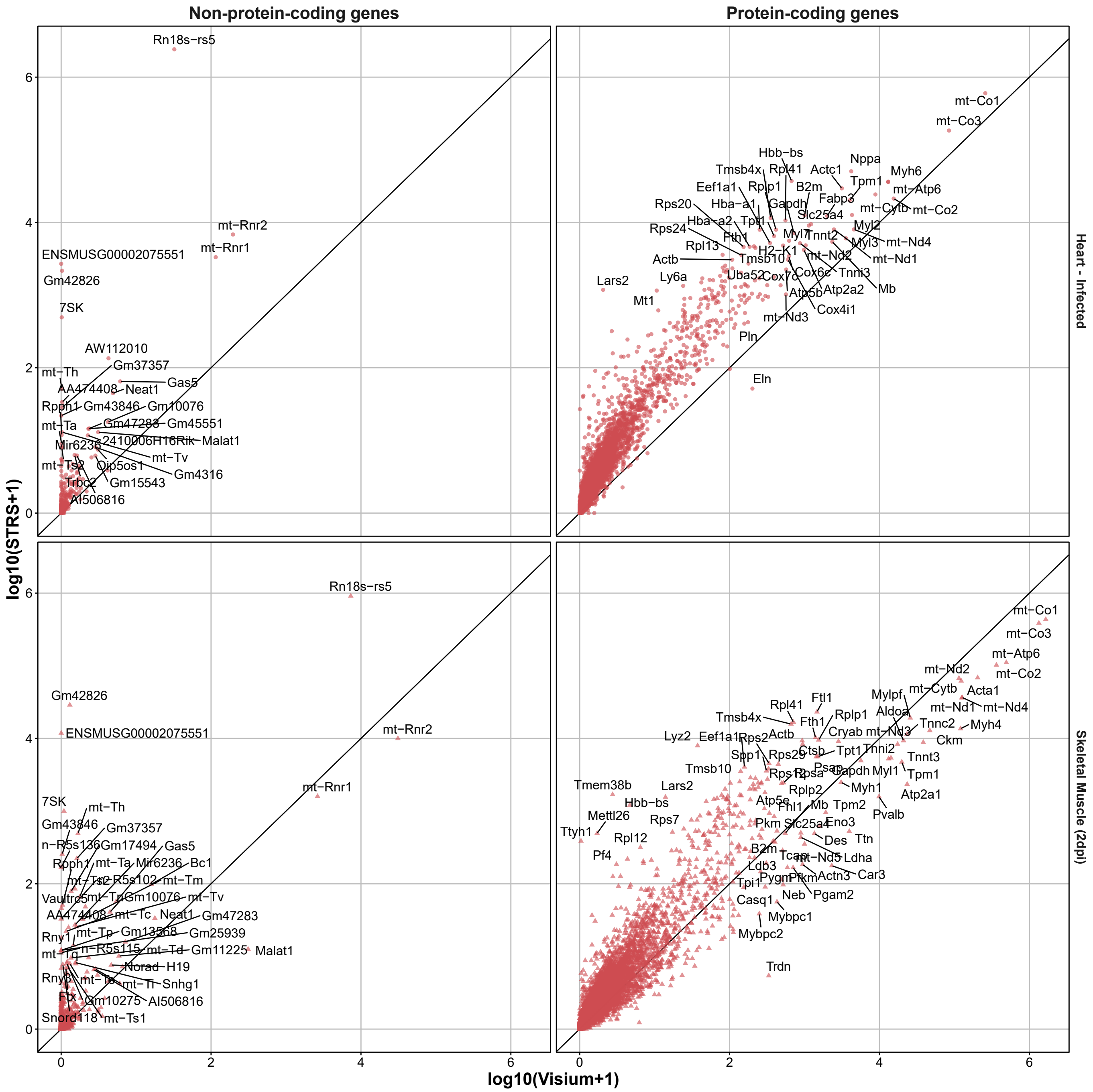


**Figure S2.** Gene-by-gene comparison across Visium and Spatial Total RNA-Sequencing. Genes are split between protein coding (right) and non-protein-coding (left) genes. Data is shown for injured skeletal muscle (2 days post-injury) and infected heart samples (Fig. 1).

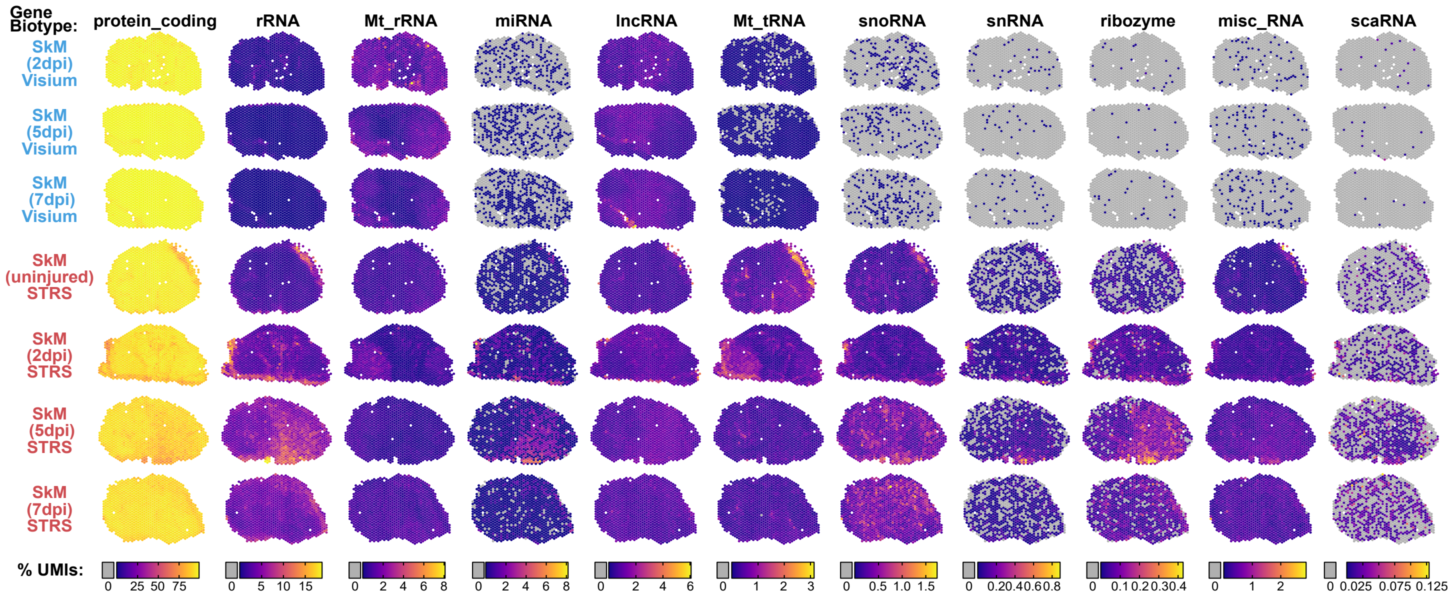


**Figure S3.** Transcript biotype spatial distribution comparison between Visium and Spatial Total RNA-Sequencing (STRS) for regenerating mouse skeletal muscle. Color scale shows the percent of unique molecules (UMIs) for each spot that correspond to each transcript biotype. Gray spots contain no molecules which correspond to the given biotype. Transcript biotypes shown include protein coding, ribosomal RNA (rRNA), mitochondrial ribosomal RNA (Mt_rRNA), microRNA (miRNA), long noncoding RNAs (lncRNA), mitochondrial transfer RNAs (Mt_tRNA), small nucleolar RNA (snoRNA), small nuclear RNA (snRNA), ribozyme, miscellaneous RNA (misc_RNA), and small Cajal body-specific RNA (scaRNA).

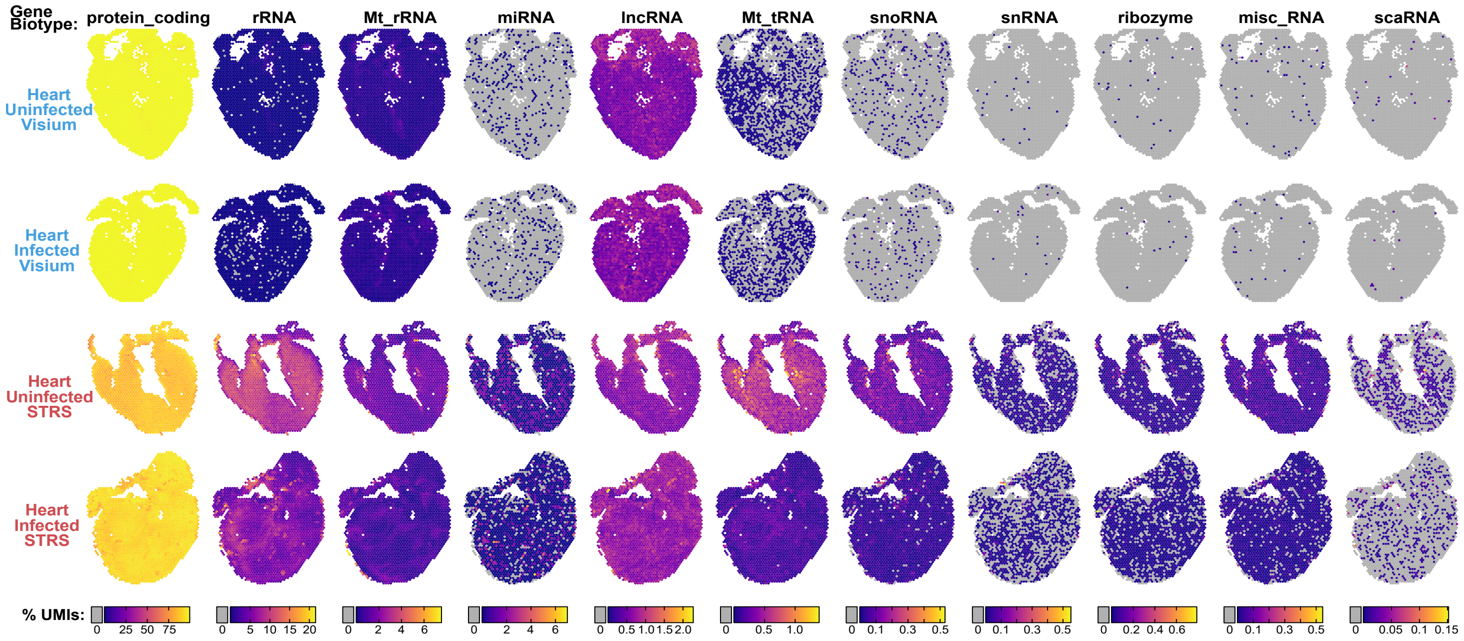


**Figure S4.** Transcript biotype spatial distribution comparison between Visium and Spatial Total RNA-Sequencing (STRS) for mouse hearts with and without Reovirus infection. Color scale shows the percent of unique molecules (UMIs) for each spot that correspond to each transcript biotype. Gray spots contain no molecules which correspond to the given biotype. Transcript biotypes shown include protein coding, ribosomal RNA (rRNA), mitochondrial ribosomal RNA (Mt_rRNA), microRNA (miRNA), long noncoding RNAs (lncRNA), mitochondrial transfer RNAs (Mt_tRNA), small nucleolar RNA (snoRNA), small nuclear RNA (snRNA), ribozyme, miscellaneous RNA (misc_RNA), and small Cajal body-specific RNA (scaRNA).

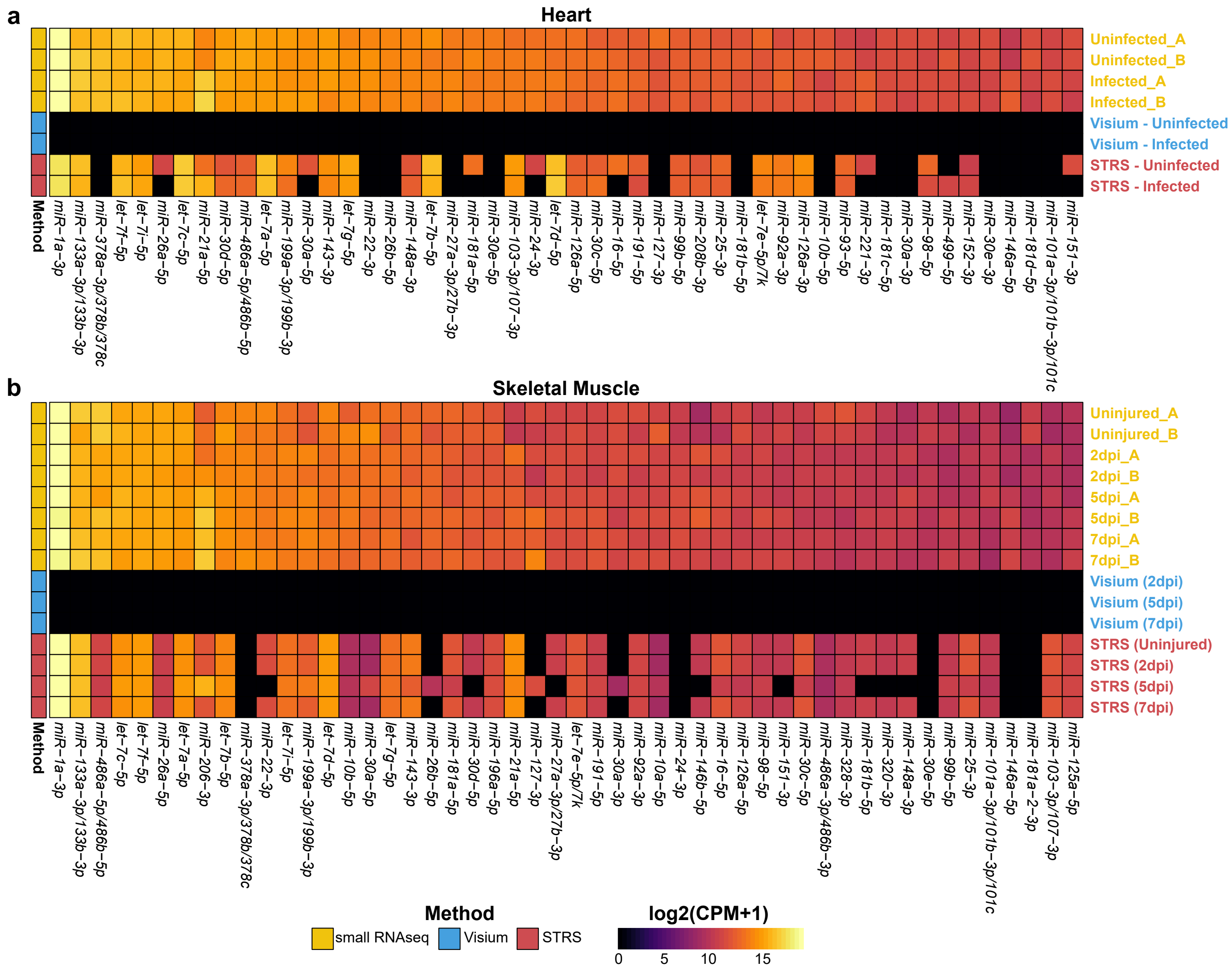


Figure S5. Comparison of mature microRNA detection in small RNA-sequencing, Visium, and Spatial Total RNA-Sequencing. Counts for (a) heart samples and (b) skeletal muscle samples are shown as log2-transformed counts per million (CPM) with a pseudocount of 1. Counts reflect UMI-deduplicated reads for STRS samples and are normalized to the total number of counts which align to mature microRNAs.

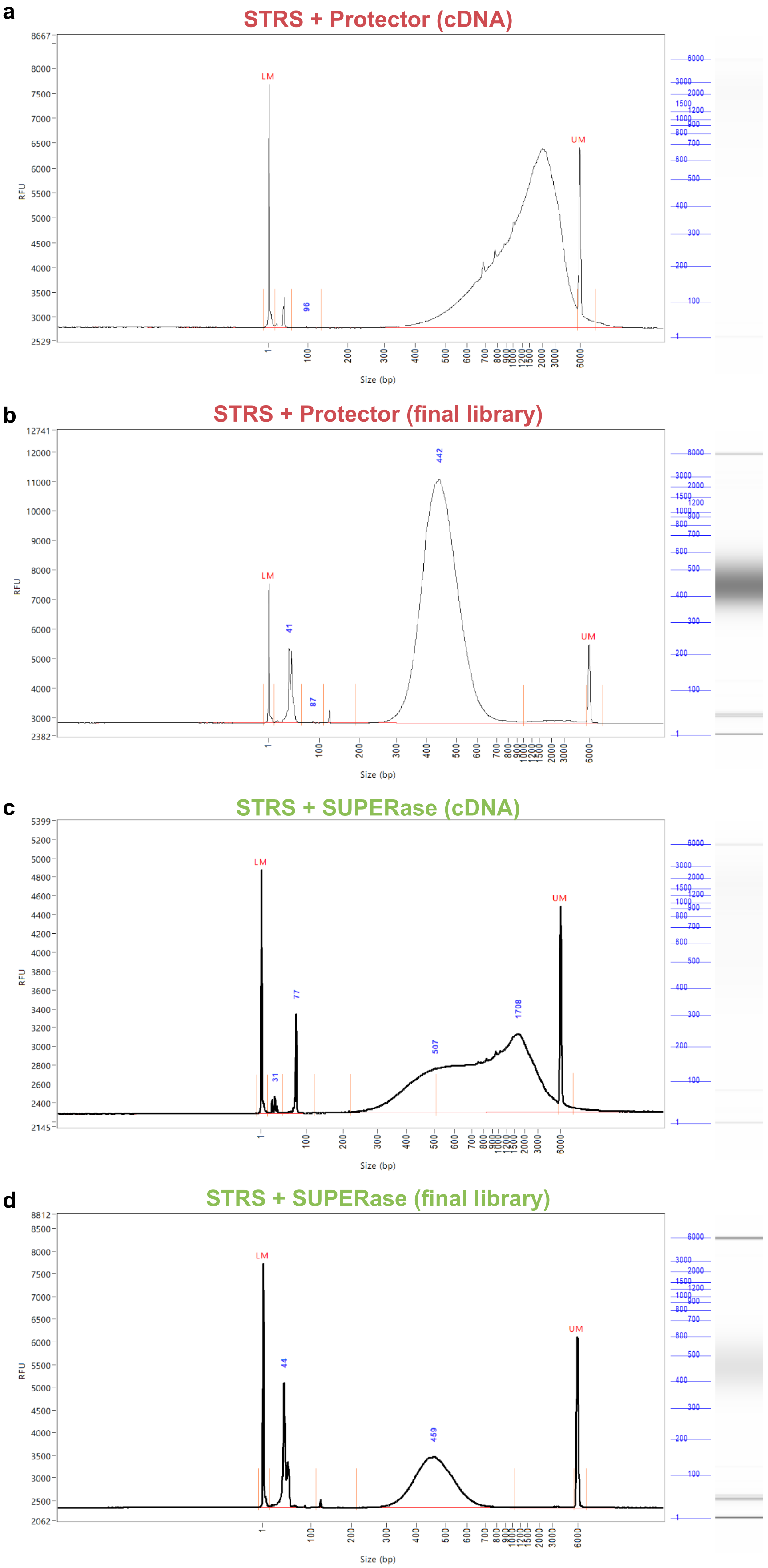


**Figure S6.** DNA fragment analysis of Spatial Total RNA-Sequencing (STRS) cDNpA (pre-tagmentation) and final libraries (post-tagmentation). Data is shown for libraries generated using either Protector RNase Inhibitor (a-b) or SUPERase in RNase Inhibitor (c-d) during in situ polyadenylation (see Methods). Samples shown are STRS_3A (GSM6034864; STRS + Protector, Uninfected Heart) and STRS_2B (GSM6034862; STRS + SUPERase, Skeletal Muscle uninjured/D0B and 2dpi).
